# BiATNovo: An Attention-based Bidirectional De Novo Sequencing Framework for Data-Independent-Acquisition Mass Spectrometry

**DOI:** 10.1101/2023.05.11.540352

**Authors:** Shu Yang, Siyu Wu, Binyang Li, Yuxiaomei Liu, Fangzheng Li, Jiaxing Qi, Qunying Wang, Xiaohui Liang, Tiannan Guo, Zhongzhi Luan

## Abstract

De novo sequencing from tandem mass spectra (MS/MS) data is a key technique for identifying novel peptides. In theory, the Data-Independent Acquisition (DIA) method can fragment all precursor ions in an unbiased and non-targeted fashion. However, each spectrum contains fragments from multiple precursor ions, and the unclear relationship between these ions and their fragments poses a significant challenge to the accuracy of de novo sequencing algorithms. Here we present BiATNovo, an attention-based bidirectional de novo peptide sequencing framework. BiATNovo comprises a bidirectional attention-based model and a bidirectional fusion-reranking post-processing module, which enables efficient capture of relationships between tandem mass spectra, fragment ions, and peptide patterns, while also expanding the candidate set to select the optimal sequence. This framework improves peptide prediction accuracy, particularly for long peptide sequences, and mitigates the imbalance where the initial amino acids are predicted more accurately than the last ones. Evaluation results demonstrate that BiATNovo outperforms existing algorithms, including DeepNovo-DIA and PepNet, in both peptid-level and amino acid-level. Furthermore, when extended to DDA datasets, BiATNovo achieves comparable performance to state-of-the-art models.

## 1 Introduction

Liquid chromatography coupled with tandem mass spectrometry (LC-MS/MS) is a fundamental technique in proteomics for protein identification within complex biological samples. Traditionally, data-dependent acquisition (DDA) has been the primary method, wherein the most abundant precursor ions are selected from an initial survey scan for subsequent fragmentation and MS/MS analysis. However, this targeted approach often leads to the exclusion of low-abundance ions, resulting in challenges such as irreproducibility and limited sampling depth.

Data-independent acquisition (DIA) [1] has emerged as a promising alternative to address these limitations. Unlike DDA, DIA fragments all precursor ions within a defined mass range during each scan, providing an unbiased and comprehensive dataset that captures information on both high and low-abundance peptides. Despite these advantages, DIA spectra present a unique challenge: each MS/MS spectrum contains a mixture of fragment ions from multiple precursor ions, obscuring the relationship between fragments and their corresponding precursors. This complexity poses significant difficulties for accurate peptide identification from DIA data, as disentangling the signals from overlapping peptide fragments remains challenging.

To address these challenges, numerous algorithms have been proposed, which can be grouped into three main categories: spectral library searching, protein sequence database searching, and de novo sequencing. Spectral library searching compares DIA spectra against pre-built libraries of known peptide fragmentation patterns, using tools such as Spectronaut [2], DIA-NN [3], and OpenSWATH [4]. Protein sequence database searching, used by tools like DIA-NN, PECAN [5], and MSFragger-DIA [6], generates theoretical spectra from protein sequence databases and matches them to DIA data. Unlike previous approaches, de novo sequencing offers a method to identify peptides directly from spectra without relying on any reference databases. This approach enables discovery of novel peptides [7], detection of post-translational modifications (PTMs), and identification of neoantigens [8].

Most de novo sequencing methods [9–13] have been developed for DDA data, while several approaches have emerged in recent years specifically targeting DIA data. One common approach utilizes tools like DIA-Umpire [14] to extract pseudo-spectra from DIA data, which are subsequently processed by DDA-based models. While this method is heavily dependent on accurate peptide extraction from MS1 spectra, posing significant challenges for low-abundance peptides, which are often overlooked. Another approach focuses on developing models tailored specifically for DIA data from the ground up. DeepNovo-DIA [15] was the first deep learning model designed for this purpose, employing a combination of convolutional neural networks (CNNs) and LSTMs to directly process DIA spectra, demonstrating that deep learning can effectively extract useful signals from complex mixed data.

Despite recent advancements, de novo sequencing accuracy for DIA data remains suboptimal. Current algorithms struggle to capture the complex relationships between spectra and peptide sequences, underscoring the necessity for more robust and specialized approaches that can address the inherent complexities of DIA data. Moreover, these models tend to perform poorly when predicting long peptide sequences. During long peptide prediction, we observed an imbalance where the initial amino acids are predicted with higher accuracy compared to those at the end. This phenomenon is common in sequencing and translation tasks, such as the prefix prediction bias observed in many neural machine translation models [16].

In this work, we present BiATNovo, an attention-based bidirectional de novo peptide sequencing framework designed to improve peptide prediction accuracy, particularly for long peptide sequences, and to address imbalance issues. BiATNovo comprises: (1) a bidirectional attention-based encoder-decoder with a local decoder, which effectively captures relationships between tandem mass spectra and fragment ions through a two-phase training strategy that learns both forward and backward sequence information; and (2) a bidirectional sequence fusion and re-ranking post-processing module, which concatenates and rescors the forward and backward peptides generated by the synchronous bidirectional model, leveraging the strengths of both to enhance sequencing accuracy and mitigate prediction imbalances. Evaluation results demonstrate that BiATNovo significantly outperforms existing de novo sequencing algorithms, such as DeepNovo-DIA and PepNet, and can be easily adapted for DDA data with minimal parameter adjustments, achieving comparable performance to state-of-the-art DDA models.

## 2 Method

In this section, we will discuss the design principles and implementation details of BiATNovo. BiATNovo is designed with the following key considerations:

### Uncovering the Relationship among tandem mass spectra, fragment ions, and sequence patterns

A key challenge in de novo peptide sequencing is identifying relationships between tandem mass spectra, fragment ions, and peptide sequences. BiATNovo addresses this by using a local decoder to extract peak features from fragment ions and a global encoder-decoder to capture broader contextual information across the entire spectrum and sequence.

### Contextual Learning

In de novo sequencing, accurate prediction of each amino acid requires a thorough understanding of its surrounding context, particularly in long peptide sequences. Each amino acid is influenced by both preceding and succeeding residues due to mass constraints and overall sequence structure. To capture this complexity, BiATNovo employs a bidirectional transformer decoder that learns dependencies in both forward and backward directions.

### Mitigating Prediction Imbalance

Current DIA peptide sequencing models often produce imbalanced results due to sequential prediction errors (described in Section 2.2). To address this, BiATNovo employs a bidirectional model to mitigate accumulated errors. In post-processing, a bidirectional fusion and reranking strategy is used to expand the peptide search space, selecting the optimal sequence from all possible candidates.

### 2.1 BiATNovo

BiATNovo framework (Fig 1) employs (1) a bidirectional sequencing model with an encoder-decoder architecture to translate MS/MS spectra into peptide sequences, and (2) a bidirectional fusion and re-ranking post-processing module to select the optimal result from generated candidates. The bidirectional sequencing model includes two components: a local decoder and an attention-based spectrum-peptide sequencing module. The output from the attention-based module is combined with a high-dimensional tensor from the local decoder and mapped to amino acid probabilities. The post-processing module then fuses the predicted amino acids from both forward and backward sequences to generate the final result. We will first introduce each component briefly, followed by a more detailed discussion in the subsequent sections.

**Fig. 1:**
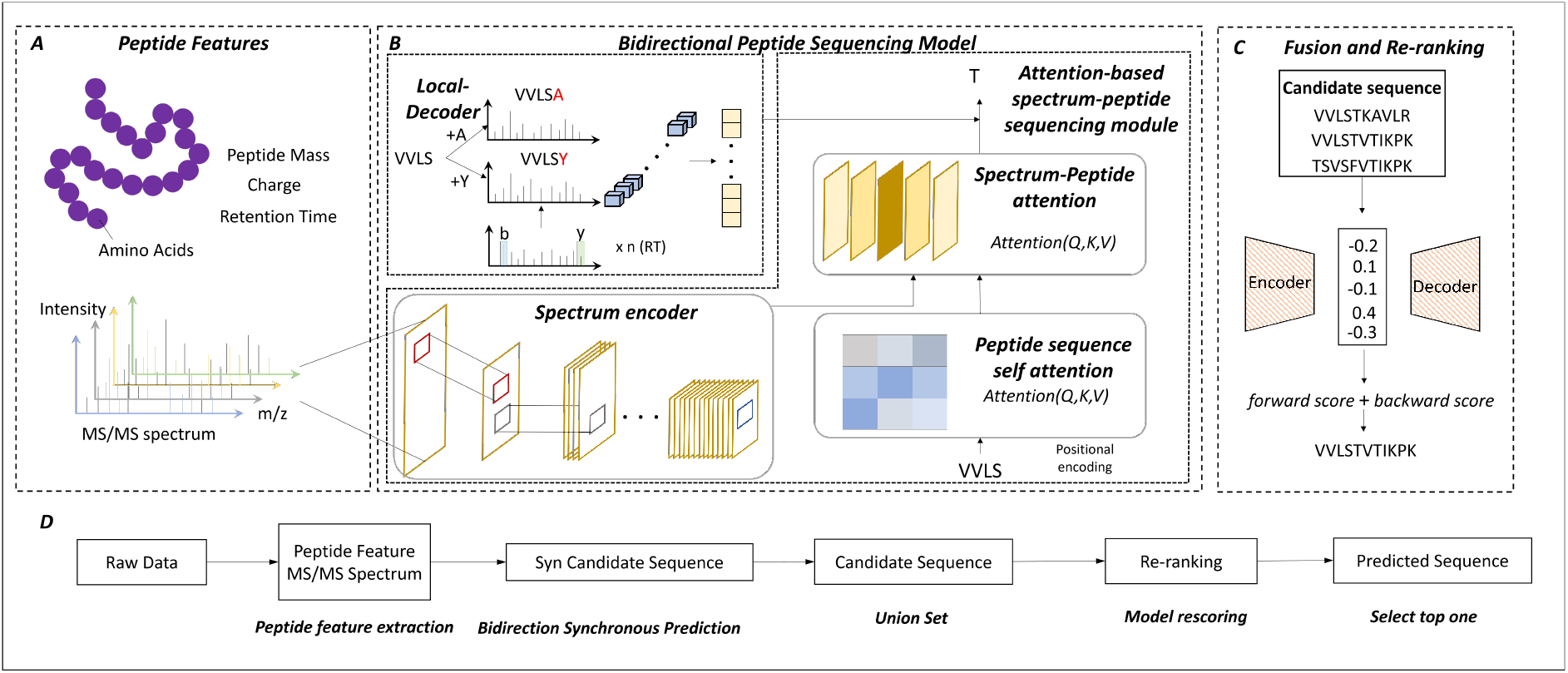
Graphical depiction of BiATNovo framework. A. represent for model feature extraction process. B. The model architecture, the local decoder and the attention-based spectrum-peptide sequencing module. C. represent for our post processing fusion and re-ranking strategy. D. The whole workflow for our inference process.

#### Bidirectional peptide sequencing model

The bidirectional peptide sequencing model consists of three key components: an attention-based spectrum-peptide sequencing module, and a local decoder. The details of bidirectional model structure will be presented at Section 2.2

##### Attention based spectrum-peptide sequencing module

This module has three main components: the Spectrum encoder, the Spectrum-Peptide Attention module and the Peptide Sequence Self-Attention module. The Spectrum encoder encodes MS/MS spectra into a high-dimensional tensor, similar to DeepNovo [9]. Multiple convolutional and pooling layers are used to derive a refined global representation of the spectra. The Spectrum-Peptide Attention module dynamically focuses on different regions of the MS/MS spectra during each step of peptide sequence prediction, assigning adaptive weights to these regions. Specifically, the Query vector is derived from the output of the self-attention [17] layer, while the Key and Value vectors are obtained from the global spectral features, enabling focused attention on key regions. The Peptide Sequence Self-Attention module captures dependencies within the peptide sequence itself. Bidirectional structure (Section 2.2) is applied across both modules, allowing them to learn contextual information in forward and backward directions, thus enhancing the model’s ability to capture long-range dependencies across the entire sequence.

##### Local decoder

The local decoder is designed to capture the correlation between theoretically possible intensity features and the MS/MS spectra at each step. Consistent with methodologies established in previous studies, such as DeepNovo [9] and DeepNovo-DIA [15].

#### Bidirectional fusion and re-ranking post-processing module

This module implements a “Fusion and Re-ranking Strategy” to reconstruct peptide sequences, producing a final result through the re-ranking of all candidate sequences, thereby improving overall prediction accuracy. Details of this strategy are provided in Section 2.3.

### 2.2 Bidirectional peptide sequencing

Peptide sequencing methods often employ single-direction decoding, predicting sequences either from left to right (forward) or right to left (backward). This is similar to text generation, where each predicted position depends on prior elements. However, single-directional decoding tends to produce imbalanced results and increased cumulative prediction errors as the sequence lengthens.

We evaluated the single-directional decoding performance of DeepNovo-DIA [15] for peptides longer than 12 amino acids on the OC, UTI, and Plasma datasets (Table 1), the results show higher accuracy for the first four amino acids compared to the last, with forward predictions generally outperforming backward ones initially, while backward predictions slightly excel at the sequence end.

**Table 1:**
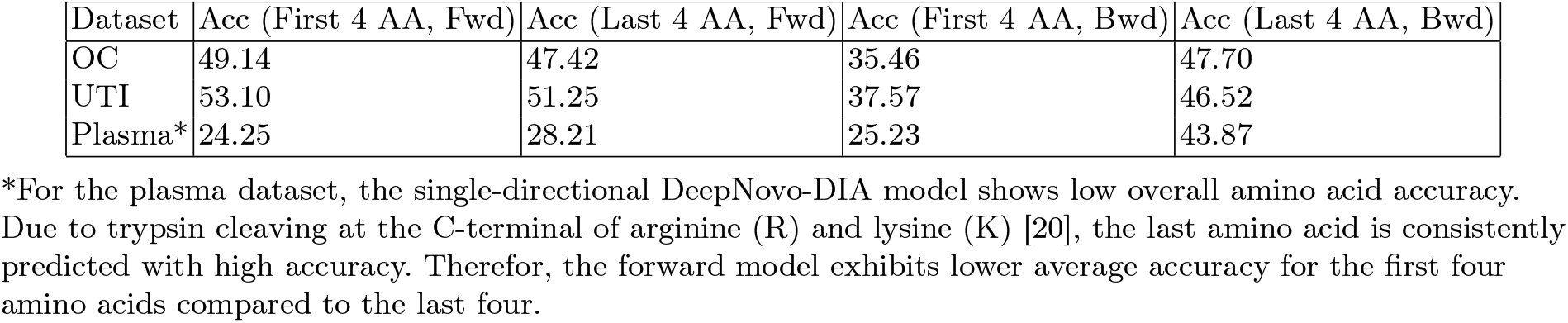
Average Prediction Accuracy for Different Peptide Sequence Positions.

| Dataset | Acc (First 4 AA, Fwd) | Acc (Last 4 AA, Fwd) | Acc (First 4 AA, Bwd) | Acc (Last 4 AA, Bwd) |
| --- | --- | --- | --- | --- |
| OC | 49.14 | 47.42 | 35.46 | 47.70 |
| UTI | 53.10 | 51.25 | 37.57 | 46.52 |
| Plasma* | 24.25 | 28.21 | 25.23 | 43.87 |

\*For the plasma dataset, the single-directional DeepNovo-DIA model shows low overall amino acid accuracy. Due to trypsin cleaving at the C-terminal of arginine (R) and lysine (K) [20], the last amino acid is consistently predicted with high accuracy. Therefore, the forward model exhibits lower average accuracy for the first four amino acids compared to the last four.

Furthermore, prediction errors tend to accumulate, affecting subsequent predictions. To tackle these challenges, we developed bidirectional peptide sequencing models to mitigate deviation accumulation and address sequence imbalance issues. By integrating bidirectional prediction, we effectively balance the use of both forward and backward context for better overall performance.

### Bidirectional independent peptide sequencing

The bidirectional independent prediction approach involves generating predictions independently from both forward and backward directions, utilizing a transformer decoder for each direction.

### Bidirectional synchronous peptide sequencing

Inspired by natural language processing techniques [18, 19], bidirectional synchronous prediction enables better convergence of forward and backward predictions at intermediate positions. Unlike independent prediction, which relies on previously predicted amino acids, bidirectional synchronous prediction incorporates both past and future context, improving peptide accuracy. Fig 2, illustrating how forward and backward predictions converge at intermediate positions to enhance the accuracy of candidate peptide sequences. When predicting amino acid ***I***, the model uses information from forward-predicted ***A*** and ***S***, as well as backward-predicted ***Q*** and ***V***, allowing for a more comprehensive use of global context.

**Fig. 2:**
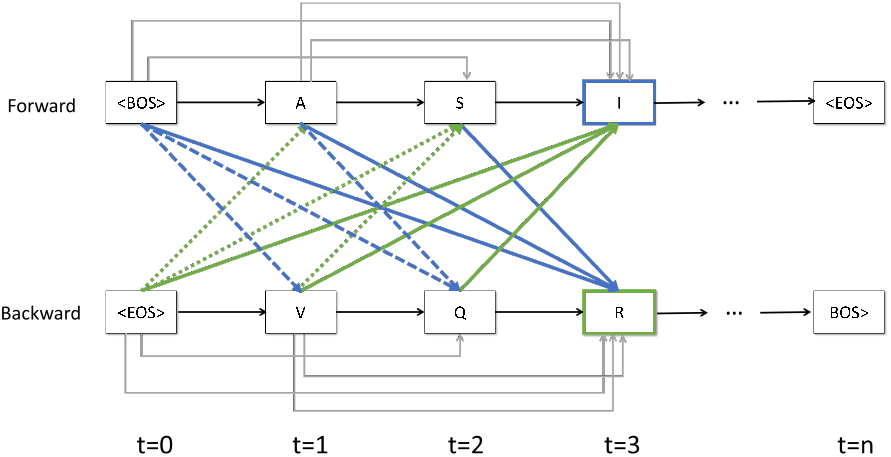
Illustration of the decoding process in the synchronous bidirectional self-attention model. Forward denotes left-to-right decoding guided by the start token ⟨BOS⟩ and backward means right-to-left decoding indicated by the start token ⟨EOS⟩. t represents the current time step in the sequence.

To achieve bidirectional synchronous prediction, we transformed the Peptide Sequence Self-Attention module into a bidirectional synchronous self-attention module. The model employs a decoder to perform left-to-right and right-to-left predictions simultaneously. This involves using bidirectional tensors for *Query, Key*, and *Value*, allowing interactions between forward and backward calculations. The resulting hidden layer vectors are merged and passed through a linear layer as input for the next layer. The computation process is illustrated in Fig 3.

**Fig. 3:**
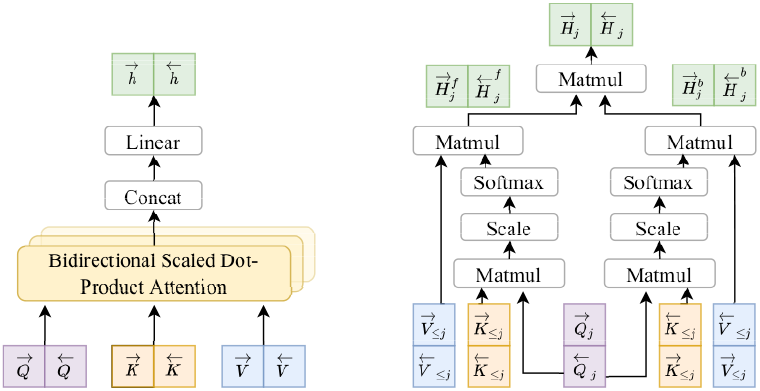
Synchronous bidirectional attention model based on scaled dot-product attention. It operates on forward and backward queries Q, keys K, values V.

In this model, *Q*_≤*j*_, *K*_≤*j*_, and *V*_≤*j*_ represent the Query, Key, and Value vectors for both forward(→) and backward(←) directions. Specifically, 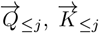, and 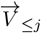 are used in forward prediction, while 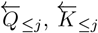 and 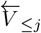 are applied in backward prediction. The corresponding hidden states are denoted as 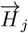 and 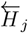.

Hidden layer outputs are decomposed into historical 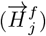 and future 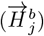 representations. Forward hidden outputs utilize past information, while backward hidden outputs incorporate future context. The attention mechanism is consistent for both directions, as defined in the equation below:

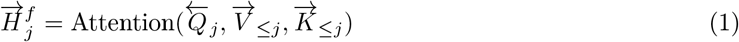

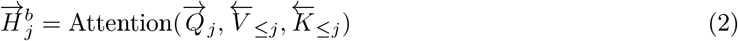

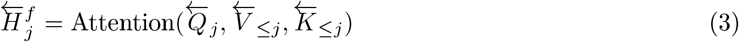

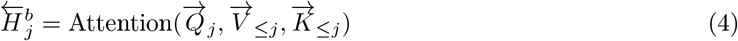

The final hidden state *H*_*j*_ for both forward and backward directions is obtained by combining the historical and future representations as follows:

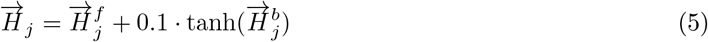

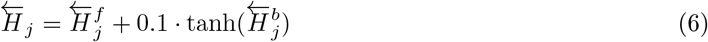

During inference, dynamic beam search is applied to facilitate bidirectional decoding. Unlike unidirectional decoding, where the beam search size is *k*, bidirectional decoding requires splitting the beam search size equally between the forward and backward directions, resulting in a beam size of 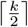 per direction.

### 2.3 Bidirectional Fusion and re-ranking Strategy

After generating candidate sequences from forward and backward predictions, the bidirectional fusion and re-ranking strategy selects the sequence with the highest score. Let the forward prediction be forward_peptide_ = {*a*_1_, *a*_2_, *a*_3_, …, *a*_*m*_} and the backward prediction be backward_peptide_ = {*b*_1_, *b*_2_, *b*_3_, …, *b*_*n*_}. Forward and backward predictions are concatenated to form a new candidate, as shown in Formula 7. To ensure validity, each candidate must meet the mass error constraints detailed in Formula 8. Where 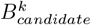 represents the *k*^*th*^ candidate sequence, while *Δ* denotes the absolute mass deviation.

All candidates are rescored using Formula 9. The final score is obtained by summing both forward and backward prediction scores, and the peptide sequence with the highest score is selected. These scores are computed by aggregating the prediction scores at each position, derived from the log-softmax probability of the predicted result at each position. A detailed calculation for the forward direction is provided in Formula 10 where *x*_*i*_ represents the amino acid at the *i*^*th*^ position of the 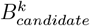 peptide sequence. The backward direction follows a similar approach. The self-attention-based bidirectional peptide sequencing method is outlined in Algorithm 1.

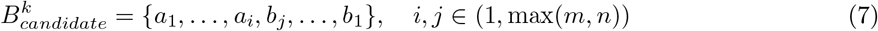

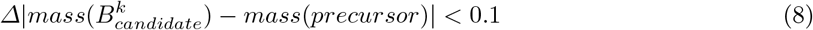

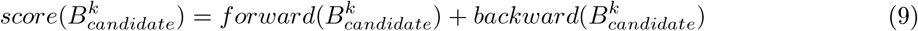

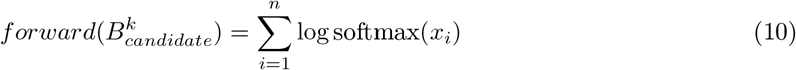

#### Algorithm 1

Bidirectional Peptide Sequencing

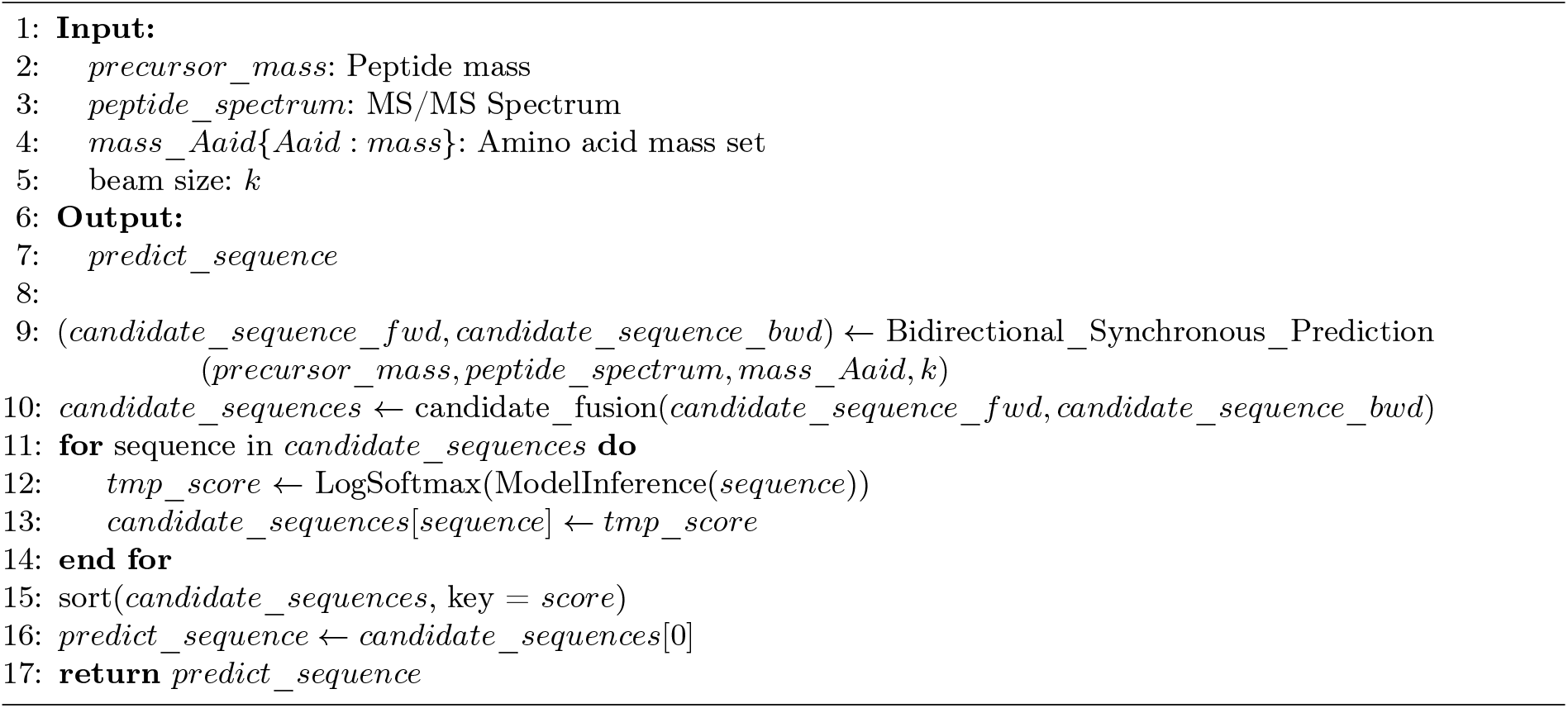

### 2.4 Training Strategy

#### Two-phases Training

The model training process consists of two distinct phases: pretraining and fine-tuning.

#### Pretraining

In this phase, the model is trained using a bidirectional independent self-attention structure with the full training dataset.

#### Fine-Tuning

In this phase, the pretrained model is used to predict peptide sequences for the entire training dataset. Subsequently, a new dataset consisting of ⟨original_feature, predicted_sequence⟩ pairs is constructed and utilized for further training. Finally, model training resumes with this new dataset, enabling the bidirectional synchronized self-attention structure.

This two-phase training strategy ensures that synchronous bidirectional decoding allows the latter half of the sequence to incorporate information from its reverse direction, improving model coherence. To maintain consistency between training and testing, pseudo peptide sequences are generated.

### 2.5 Code availability

Our code and model weights are available at BiATNovo-DIA via the MIT license (Copyright (c) 2024 Shu Yang). We also provide the code and model weights for DDA dataset at BiATNovo-DDA under the MIT license (Copyright (c) 2024 Shu Yang)

## 3 Results

### 3.1 Datasets and experiments setting

Following the evaluation method established by DeepNovo-DIA, we used the same original DIA dataset, which includes both training and test sets. The raw files for this dataset are available from the original publications [21, 22]. In de novo sequencing experiments, peptides identified through database searches are typically used as ground truth. To create our training and test datasets, we used DIA-NN, a widely used database search tool, and applied strict filtering criteria to keep only those entries with a false discovery rate (FDR) of 1% or lower and a precursor mass error within 10 ppm as the ground truth. The training set consists of urine samples from 64 subjects, comprising 2,177,667 spectra and 553,828 labeled features. We split the dataset into training and validation subsets in a 9:1 ratio, ensuring no peptide overlap to maintain data integrity and evaluate model generalization.

Due to the high-dimensional and complex nature of DIA datasets, they are particularly susceptible to overfitting. In the first phase of training, we trained the independent bidirectional model for six epochs. In the second phase, we fine-tuned the synchronous bidirectional model for an additional two epochs, using early stopping to save the best-performing model based on validation set performance. We used the Adam optimizer with a dynamic learning rate schedule, starting with a higher rate and gradually decreasing it based on validation loss to achieve faster initial convergence and finer adjustments as training progressed. For evaluation, we used two additional datasets: ovarian cyst (OC) and urinary tract infection (UTI) subjects (six subjects each), along with a plasma sample dataset [22]. These test datasets were excluded from model development to ensure unbiased performance assessment.

To assess the performance of BiATNovo on DDA and benchmark it against state-of-the-art methods such as DeepNovo [9] and CasaNovo [11, 13], we used the nine-species benchmark dataset and evaluation framework initially introduced by Tran et al. (2017) [9], which has been utilized in several subsequent studies. In this experiment, we adopted a leave-one-out cross-validation strategy. Specifically, eight species datasets were used for training and validation, while one species dataset was held out for testing. The training and validation sets were divided in a 9:1 ratio, after which the model was evaluated on the held-out species dataset. Each training session on the eight-species datasets ran for 30 epochs.

### 3.2 Evaluation metrics

We employ evaluation metrics consistent with prior research to benchmark BiATNovo’s de novo peptide sequencing predictions against ground-truth sequences obtained through database searches. Specifically, our evaluation is conducted at both the amino acid and peptide levels: amino acid-level accuracy is defined as the proportion of correctly predicted amino acids out of the total amino acids, while peptide-level accuracy represents the proportion of fully correct peptide predictions relative to the total number of predicted peptides. Additionally, we use a precision-coverage curve to assess model performance, illustrating the trade-off between prediction accuracy (precision) and the proportion of peptides for which predictions are successfully generated (coverage).

### 3.3 Performance Evaluation

#### Overall Comparative Analysis of BiATNovo and State-of-the-Art (SOAT) Models

To ensure a fair comparison, both DeepNovo-DIA and BiATNovo were trained and tested on the same dataset. DeepNovo-DIA was retrained with its original configuration, but using our training data generated by DIA-NN. Our results, in Table 2 show significant improvement on the OC and UTI datasets, with BiATNovo achieving an average increase of 10% in peptide-level accuracy and 12% in amino acid-level accuracy compared to DeepNovo-DIA. The precision-coverage subplots in Figure 4 further demonstrate that BiATNovo consistently outperforms DeepNovo-DIA across all coverage levels. Additionally, PepNet [12], a model trained on DDA-derived MS/MS spectra, was included for further comparison due to its strong performance on plasma datasets. As shown in Table 2, BiATNovo improves peptide-level precision by 3% compared to DeepNovo-DIA and by 6% compared to PepNet. The precision-coverage curve in Figure 4 also shows that BiATNovo consistently outperforms both models on the plasma dataset. The results on the test set indicate that our bidirectional framework is more effective in capturing the relationship between highly multiplexed spectra and peptides. Additionally, the accuracy on the plasma dataset could potentially be improved through fine-tuning. We provide further discussion on this point in Section 4.

**Table 2:** Comparison of DeepNovo-DIA, PepNet and BiATNovo.

| Evaluation | OC Dataset |  | UTI Dataset |  | Plasma Dataset |  |  |
| --- | --- | --- | --- | --- | --- | --- | --- |
|  | DeepNovo-DIA | BiATNovo | DeepNovo-DIA | BiATNovo | DeepNovo-DIA | PepNet | BiATNovo |
| Amino acid-level precision(%) | 55.62 | 68.17 | 62.86 | 71.08 | 43.54 | 40.08 | 45.25 |
| Amino acid-level recall(%) | 55.96 | 67.46 | 62.43 | 70.76 | 43.35 | 40.04 | 45.11 |
| Peptide recall number | 31153 | 39344 | 21725 | 25884 | 2755 | 2951 | 3277 |
| Peptide-level precision(%) | 50.98 | 64.39 | 55.25 | 65.84 | 33.22 | 30.64 | 36.91 |

**Fig. 4:**
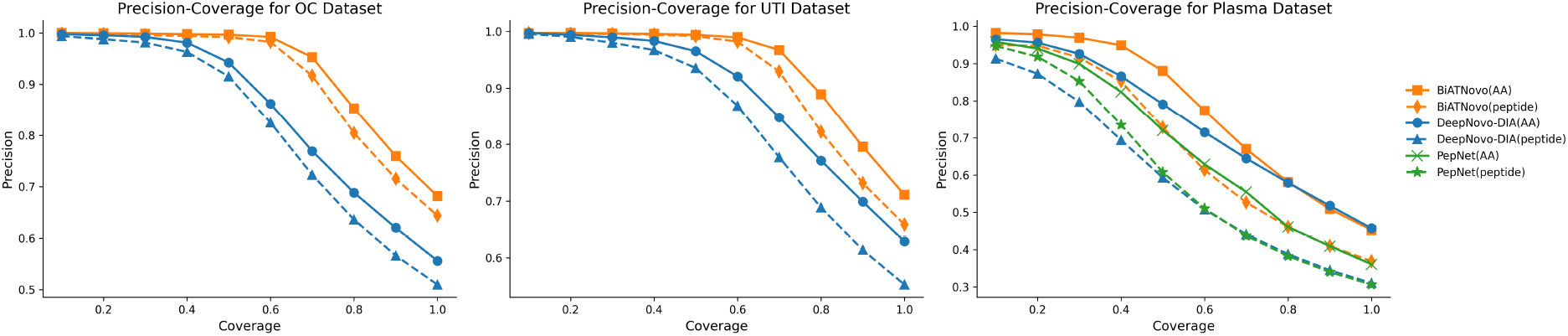
Precision-coverage curves for BiATNovo and SOTA Models. The curves are generated by ranking the predicted peptides based on their confidence scores. For the amino acid-level curves, every amino acid within a peptide is assigned the same score as the peptide itself.

#### Evaluation of Long Peptide Prediction Performance and Imbalance Issues

As illustrated in Figure 5, both DeepNovo-DIA and PepNet exhibit a noticeable decline in performance as peptide length increases, particularly when the peptide length exceeds 18, where peptide-level accuracy drops by more than half. This decrease is likely due to the increased complexity of MS/MS spectra for long sequences and the accumulation of decoding errors during prediction.

**Fig. 5:**
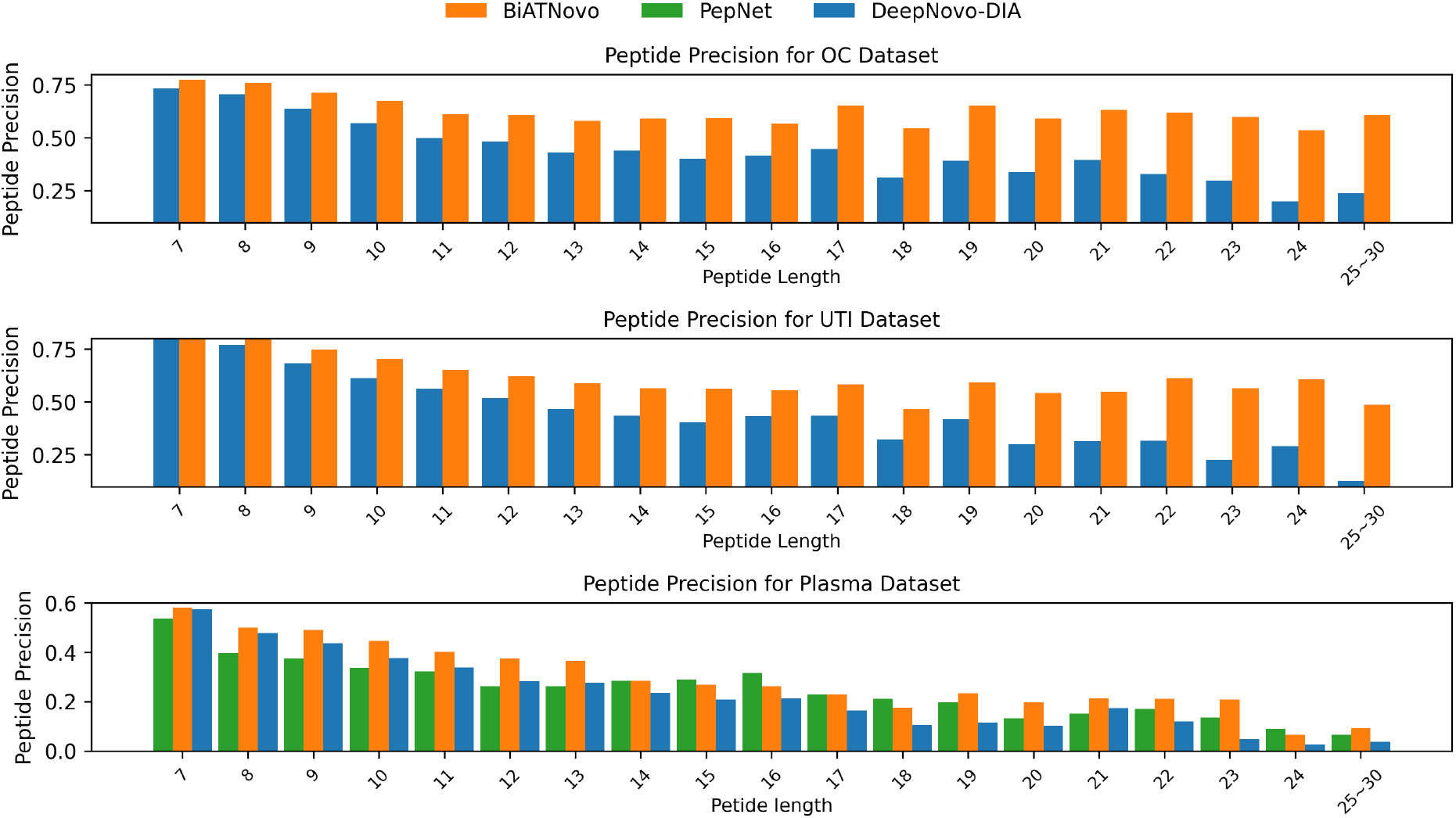
Peptide-level precision across different peptide lengths. Peptides were grouped as follows: lengths greater than 25 formed a group, and the remaining lengths were grouped individually by unit length.

In the plasma dataset, BiATNovo consistently achieves higher peptide precision compared to the other models across nearly all peptide lengths. This trend is even more pronounced in the OC and UTI datasets, where BiATNovo’s performance remains stable with increasing peptide length, showing minimal decline. These results indicate that BiATNovo is well-suited for handling long peptides and performs robustly in more challenging scenarios.

To quantify the imbalance level in a model, we introduced a metric called difference−*i* which measures the absolute difference between the accuracy at position *i* on the forward and backward sides. Table 3 illustrates these results. From the table, we can observe that BiATNovo notably mitigates the imbalance issue compared to DeepNovo-DIA.

**Table 3:** Difference-i Metric Evaluation for DeepNovo-DIA and BiATNovo.

| Evaluation | OC |  | UTI |  | Plasma |  |
| --- | --- | --- | --- | --- | --- | --- |
|  | DeepNovo-DIA | BiATNovo | DeepNovo-DIA | BiATNovo | DeepNovo-DIA | BiATNovo |
| 25th Percentile (Q1) | 2.192 | 1.581 | 4.205 | 3.228 | 3.086 | 2.118 |
| Median (Q2) | 7.295 | 4.429 | 7.973 | 4.614 | 15.340 | 11.988 |
| 75th Percentile (Q3) | 10.222 | 6.836 | 13.689 | 6.947 | 20.000 | 17.416 |

The improvement in handling long peptide sequences and reducing prediction imbalance in BiATNovo comes from its two parts: (1) In the bidirectional synchronous peptide sequencing, each amino acid (AA) prediction incorporates information from both the forward and backward directions, unlike DeepNovo-DIA and other left-to-right or right-to-left models, which only consider information from a single direction. This bidirectional approach provides richer context for predicting each AA, resulting in improved accuracy and a reduction in prediction imbalance. (2) sequence fusion and re-ranking strategy combines and re-scores the forward and backward peptide sequences generated by the bidirectional model. This method takes advantage of the more accurate segments from both directions, enhancing sequencing accuracy.

#### Ablation Experiments

To assess the performance of different components within the BiATNovo framework, we evaluated three variants. The first variant (BiATNovo base) utilizes a bidirectional attention-based model with an independent self-attention network. The second variant (BiATNovo w sync), fine-tuned from the first, incorporates a bidirectional synchronous structure. The third variant (BiATNovo final) further integrates a bidirectional fusion-reranking post-processing module. As shown in Table 4, the bidirectional synchronous network achieves a 7% and 9% improvement in amino acid-level and peptide-level precision, respectively, over the bidirectional independent network, demonstrating the advantage of leveraging bidirectional information for enhancing peptide sequencing accuracy. Moreover, the area under the curve (AUC) metric indicates an additional 2% improvement when the bidirectional fusion-reranking module is employed.

**Table 4:** Performance Comparison of Different Variants of BiATNovo on OC Dataset.

| Variants | Amino Acid |  |  | Peptide |  |  |
| --- | --- | --- | --- | --- | --- | --- |
|  | Precision | AUC | AUC-Increment | Precision | AUC | AUC-Increment |
| BiATNovo base | 0.603 | 0.768 | — | 0.542 | 0.744 | — |
| BiATNovo w sync | 0.672 | 0.813 | 0.044 | 0.632 | 0.801 | 0.05 |
| BiATNovo final | 0.681 | 0.839 | 0.07 | 0.643 | 0.822 | 0.081 |

#### Performance Of BiATNovo On Data-Dependent acquisition (DDA) Data

We evaluated BiATNovo against the current state-of-the-art DDA models, CasaNovo and DeepNovo. As shown in Figure 6, BiATNovo and DeepNovo-DIA both outperform CasaNovo significantly in terms of amino acid-level accuracy. BiATNovo achieved the highest amino acid-level precision. The superior performance compared to CasaNovo may be attributed to the use of a local decoder. In each prediction step, the local decoder captures specific peaks signal, which enhances amino acid-level accuracy.

**Fig. 6:**
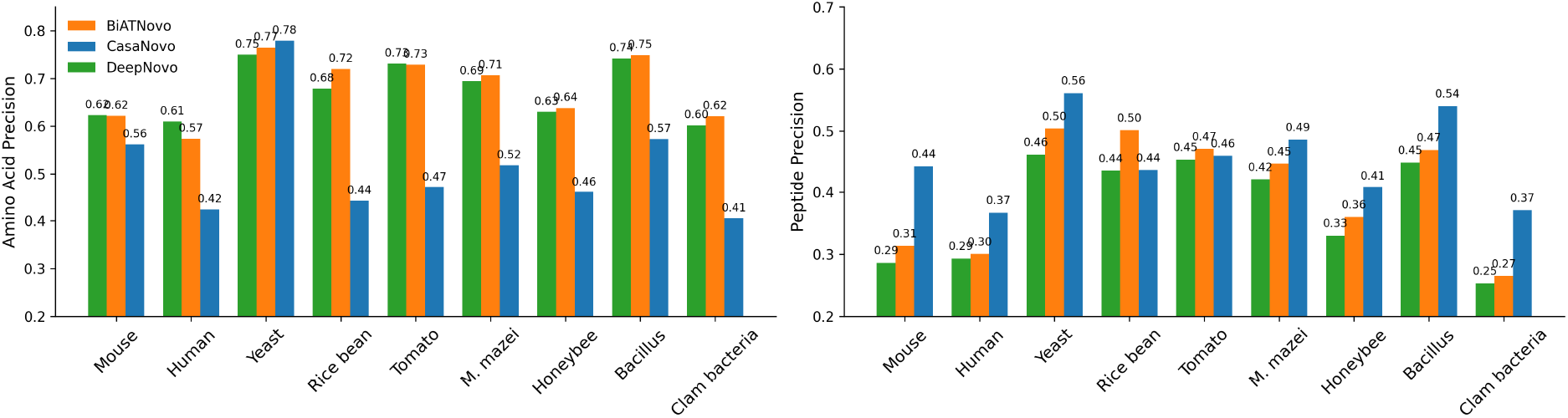
Performance comparison of BiATNovo on DDA datasets against DeepNovo and CasaNovo, in terms of Amino Acid-Level Precision and Peptide-Level Precision.

For peptide-level precision, BiATNovo surpasses the DDA model in the species Rice Bean and Tomato. In other species, CasaNovo achieves higher peptide-level accuracy, likely due to its global transformer encoding approach, which may better capture global signals and enhance peptid-level precision.

## 4 Discussion

In this paper, we present an attention-based bidirectional de novo sequencing framework specifically designed for Data-Independent Acquisition (DIA) data. The framework demonstrates strong performance even with long peptide sequences and addresses the imbalance issue in peptide sequencing through a bidirectional attention-based model and a bidirectional fusion-reranking post-processing module. In our dataset evaluation, the model achieves up to a 10% improvement in amino acid accuracy and up to a 14% improvement in peptide-level accuracy over DeepNovo-DIA.

The relatively reduced accuracy on the Plasma dataset compared to other datasets, is likely due to the significant distribution differences, as our training set consists of urine samples. To further investigate, we randomly selected 1,000 peptide sequences from the plasma test data after shuffling and fine-tuned BiATNovo. This fine-tuning resulted in a 12% improvement in both peptide and amino acid accuracy for plasma data, suggesting that our model can be used as a pre-trained model. In the future, we plan to provide pre-training models for multiple tissue samples, enabling users to fine-tune the models with minimal data for de novo sequencing of different tissue samples.

In the evaluation of nine species on DDA datasets, we found that our model achieved significantly higher amino acid accuracy compared to Casanovo. However, Casanovo exhibited better peptide-level accuracy across a wider range of species. This discrepancy might be due to our model’s use of a local decoder, which captures stronger local signals (candidate ion peaks) at each prediction step, thereby enhancing amino acid accuracy. In contrast, Casanovo employs a transformer to encode the global spectrum, which allows it to learn and recover global patterns more effectively, especially when spectra contain noise or missing ions, leading to improved peptide-level accuracy. Moving forward, we plan to integrate a transformer encoder to model the global spectrum, as this may further boost peptide-level accuracy.

